# Phylodynamic analysis of the historical spread of Toscana virus around the Mediterranean

**DOI:** 10.1101/380477

**Authors:** M. Grazia Cusi, Claudia Gandolfo, Gianni Gori Savellini, Chiara Terrosi, Rebecca A. Sadler, Derek Gatherer

## Abstract

All available sequences of the three genome segments of Toscana virus with date and location of sampling were analysed using Bayesian phylodynamic methods. We estimate that extant Toscana virus strains had a common ancestor in the late 16^th^ to early 17^th^ century AD, in territories controlled by the Ottoman Empire, giving rise to an ancestral genotype A/B in north Africa and to genotype C in the Balkans. Subsequent spread into western Europe may have occurred during the period of European colonization of north Africa in the 19^th^ and early 20^th^ centuries AD, establishing genotypes A and B in Italy and Spain respectively. Very little positive evolutionary selection pressure is detectable in Toscana virus, suggesting that the virus has become well adapted to its human hosts. There is also no convincing evidence of reassortment between genome segments, despite genotypes A and B now co-circulating in several countries.

## 1. Introduction

The International Committee on Taxonomy of Viruses (ICTV) defines Toscana virus (TosV) as a member of the species *Sandfly fever Naples phlebovirus* (Order *Bunyavirales;* Family *Phenuiviridae;* Genus *Phlebovirus)* (ICTV, 2016). TosV is a negative-strand single-stranded segmented RNA virus with three genome segments of 6.4 kb, 4.2 kb and 1.9 kb, designated L, M and S respectively. The L segment (Accardi et al., 1993) encodes the RNA-dependent RNA polymerase. The M segment encodes a polyprotein which is the precursor of G1 and G2 glycoproteins and of a non-structural protein, NS_M_ (Di Bonito et al., 1997). The S segment encodes two proteins in complementary orientation, the nucleoprotein (N) and the non-structural (NS) protein (Giorgi et al., 1991).

Sandfly fever is transmitted to humans by the bite of sandflies of the genus *Phlebotomus* and was first described in Egypt in 1910 (Wakeling, 1910). Various viruses can cause it (Alkan et al., 2013; Depaquit et al., 2010), and TosV was first isolated in Italy in the early 1980s (Verani et al., 1984). Although TosV infection is usually self-limiting, a minority of cases develop meningitis or meningoencephalitis (Nicoletti et al., 1991), accounting for 95% of aseptic meningitis cases occurring in Mediterranean countries during the summer (Cusi et al., 2010). Sandfly fever caused by TosV is therefore southern Europe’s most significant autochthonous arboviral disease of humans.

Phylogenetic analysis of the M genome segment showed that the Italian strains of TosV can be classified into four lineages G1 to G4 (Venturi et al., 2007). Analysis of a set of S segment nucleoprotein genes from Italian, French and Spanish strains demonstrated the presence of two broader genotypes (Charrel et al., 2007; Sanbonmatsu-Gamez et al., 2005), later designated A and B (Collao et al., 2009) with lineages G1 to G4 forming sub-genotypes of A. The division into genotypes A and B is found in all three segments (Collao et al., 2009), suggesting geographical isolation separating predominantly Italian strains (genotype A) from most French and all Spanish strains (genotype B) and no reassortment of segments between genotypes. Bayesian phylogenetic analysis (Zehender et al., 2009) of the time of the most recent common ancestor (tMRCA) of the M segment of TosV genotype A, gave 376 years before present (yBP), but with wide confidence limits – in Bayesian terminology, 95% highest posterior density (HPD) ranges – of 75 to 856 yBP. A Bayesian skyline plot (Zehender et al., 2009) showed that there had been a decrease in the number of TosV infections since the 1970s, consistent with observations of decreasing population TosV seropositivity over time (de Ory-Manchon et al., 2007) and greater frequency of seropositivity in older individuals (de Ory-Manchon et al., 2007; Terrosi et al., 2009).

A strain of TosV was subsequently isolated in Croatia (Punda-Polic et al., 2012) with S and L segments that both appeared to be outgroups of the defined genotypes A and B. Together with an L segment isolated in Greece (Papa et al., 2014), this defines genotype C. The presence of the TosV genotype C outgroup in the Balkans, but not in the western Mediterranean, initially suggests a westward movement of the virus which, based on previous Bayesian phylogenetic analyses (Collao et al., 2009; Zehender et al., 2009), has occurred over a timescale of hundreds to a few thousand years.

In this paper, we seek to clarify the evolutionary history of Toscana virus by consolidating all available sequence data, for all three segments, which is suitable for Bayesian phylodynamic analysis. We perform both phylogeographic analysis to establish the direction of diffusion of the virus and phylogenetic analysis to establish the timescale of that diffusion. For these analyses, the segments must respectively have a site or time of sampling specified. Our goal is to assess the evidence for both the direction and time of spread of TosV across the Mediterranean. Furthermore, we also analysed all three segments for evidence of natural selection. A previous study suggested that amino acid residue 338 from the M segment polyprotein is under positive selection, but that the predominant mode of molecular evolution was constraint with the ratio of non-synonymous to synonymous substitution rates (dN/dS, or omega) at approximately 0.18 (Venturi et al., 2007).

## 2. Methods

### 2.1 Collection of Phlebovirus genome segment sequences for maximum parsimony phylogenetic analysis

To place TosV in the context of the genus *Phlebovirus* as a whole, we downloaded all nucleotide sequences classified by GenBank as from that genus. In July 2017, 3741 such sequences were available. Since the taxonomy of *Phlebovirus* is complex, we assumed that each virus with a different name was potentially a representative of a different species. Sequences annotated as partial were removed, except in cases where only a partial sequence was available for any potential (i.e. uniquely named) species. This produced L, M and S sets of 93, 82 and 103 sequences respectively. These were aligned using MAFFT (Katoh and Standley, 2014). For the L and M segments, 5’ and 3’ untranslated sequences were removed and the alignment was refined using Muscle (Edgar, 2004) in “align codons” mode within MEGA (Kumar et al., 2008). The best substitution model was derived in MEGA and guide neighbour joining trees were drawn (Supplementary Figures 1-3). These were used to define the outgroup species within the *Sandfly fever Naples phlebovirus* species cluster as Gordil virus. All sequences internal to Gordil virus on the guide tree were selected and realigned. A maximum parsimony tree using the max-mini branch-and-bound algorithm was then produced in MEGA for each genome segment of the *Sandfly fever Naples phlebovirus* species cluster. 100 bootstrap replicates were used to assess the reliability of the internal structure of the tree.

### 2.2 Collection of Toscana virus genome segment sequences for Bayesian phylogeographic analysis

We downloaded all nucleotide sequences classified by GenBank as TosV. In July 2017, 291 such sequences were available. Sequences with collection location were selected from the initial set of 291 TosV sequences using *tempus_et_locus* (TeL) (Carter and Gatherer, 2016). The S sequences were additionally split into nucleoprotein (N) and non-structural (NS) coding sequences (referred to subsequently as S-N and S-NS). Alignments of 100 L, 67 M, 104 S-N and 54 S-NS were produced as described in section 2.1. These are listed in Supplementary Tables 1 to 4. As TosV is a segmented virus, some strains may be reassortants. Potential reassorted strains are listed in Supplementary Table 5.

### 2.3 Collection of Toscana virus genome segment sequences for Bayesian phylogenetic analysis

Since previous Bayesian phylogenetic analyses have suffered from wide 95% HPD values, all sequences in the initial set of 291 that were less than 1 kb in length, were removed. Sequences with collection date were then selected using *tempus_et_locus* (TeL) (Carter and Gatherer, 2016). From these, alignments of 18 L, 62 M, 46 S-N and 52 S-NS were produced as described in section 2.1. These are listed in Supplementary Tables 6 to 9.

### 2.4 Parameters of Bayesian phylogenetic and phylogeographic analysis using BEAST

BEAST (Drummond et al., 2012) was run for 250 million iterations, using a GTR+G+I substitution model for the L segment and a TN93+G substitution model for the others, chosen by previous selection of the best model in MEGA. A Bayesian skyline (Drummond et al., 2005) tree model was chosen, for consistency with previous work (Collao et al., 2009; Zehender et al., 2009). The clock model was relaxed exponential, the most appropriate given the wide 95% HPDs obtained in previous studies (Collao et al., 2009; Zehender et al., 2009). Each codon position was allocated to a separate partition to estimate substitutions rates in synonymous and non-synonymous positions. Neighbour joining trees were produced in MEGA to assess clock-like behaviour in TempEST (Rambaut et al., 2016). These are downloadable from http://dx.doi.org/10.17635/lancaster/researchdata/185. In general, root-to-tip distance correlated poorly with sampling time, providing an explanation for previously published wide 95% HPDs. In an attempt to constrain the molecular clock estimates within reasonable limits, a prior was set based on substitution rates obtained from TosV previously and from other bunyaviruses. These are shown in Supplementary Table 10. Based on these, a prior of 2.7 × 10^−4^ (95% HPD 1.11× 10^−4^ to 4.24× 10^−4^ on a normal distribution) substitutions/site/year was applied to each segment. Output of BEAST was assessed using Tracer (Rambaut et al., 2014) and TreeAnnotator in the BEAST package was used to produce summary trees for each segment. A burn-in of 50 million states was used with posterior probability limit for annotation of each node set at 0.667. The topology of the Bayesian trees produced in BEAST was confirmed by bootstrap maximum likelihood trees produced in MEGA. Bayesian phylogeographic maps were superposed on a map of Europe using SPREAD v.1.0.6 (Bielejec et al., 2011) and ArcMap (http://desktop.arcgis.com/en/arcmap/).

### 2.5 Analysis of positive selection using Slr

The same alignments used for phylogeographic analysis were assessed for traces of positive selection, neutrality and constraint on a site-wise basis using Slr (Massingham and Goldman, 2005) (version 1.5.0). An initial estimate of the transition/transversion (kappa) bias was obtained in MEGA and was used along with an initial value of omega (dN/dS) of 1 (corresponding to neutral evolution) in the Slr input parameters. After the first run, the values of kappa and omega derived by Slr were used as input parameters for a second confirmatory run.

## 3. Results

### 3.1 Maximum parsimony trees of L, M and S segments in the Sandfly fever Naples phlebovirus species cluster

Based on the neighbour-joining trees for the wider genus (Supplementary Figures 1-3), the outgroup of the *Sandfly fever Naples phlebovirus* species cluster was defined as Gordil virus. Figure 1 shows a maximum parsimony tree for the L segment polymerase gene coding sequence. TosV is most closely related to the Zerdali/Tehran virus pair, with bootstrap confidence of 70%. The next outgroup is a quartet of Massilia/Granada/Arrabida/Punique viruses at boostrap of 71%. Maximum parsimony trees for the M and S segments are shown in Supplementary Figures 4 and 5, respectively.

**Figure 1.**
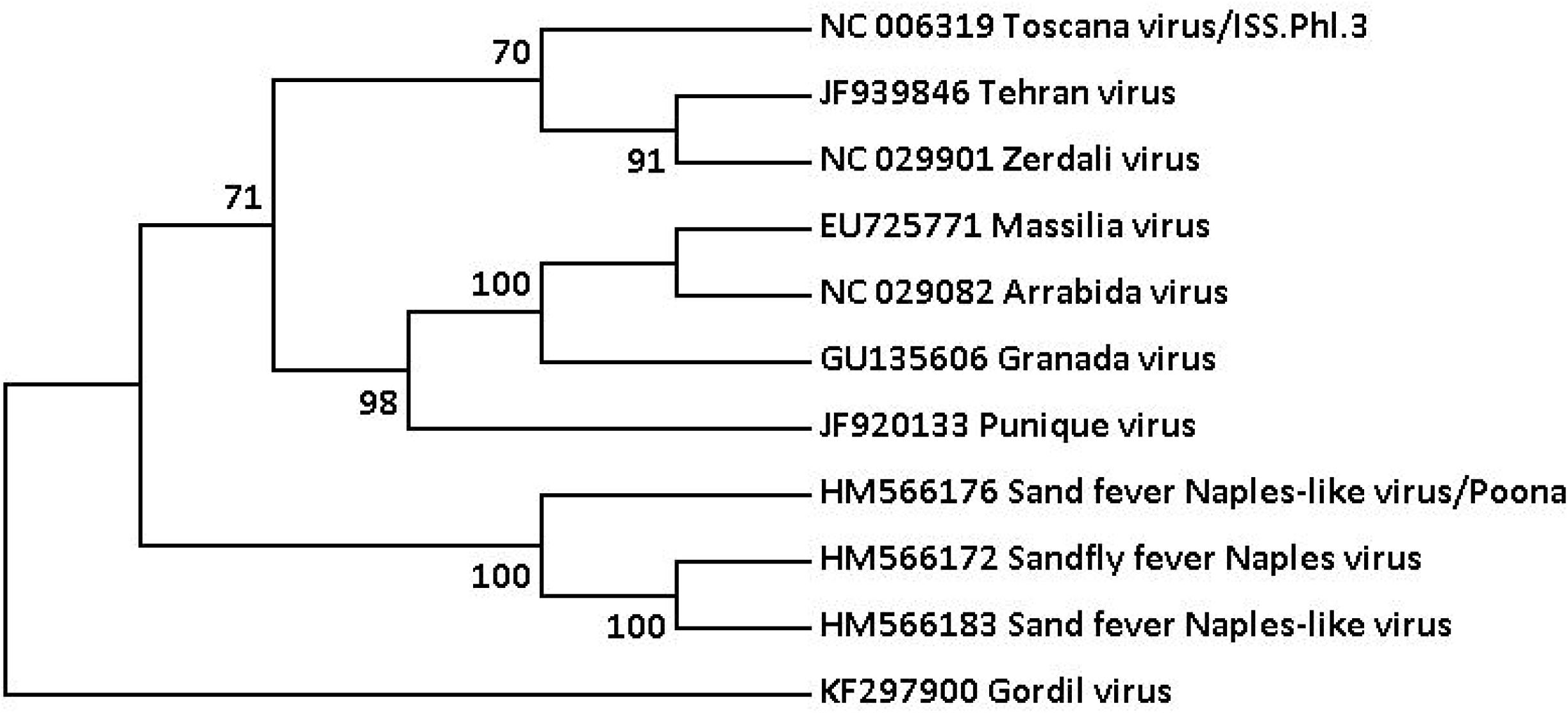
Maximum parsimony tree showing the phylogenetic position of Toscana virus within the *Sandfly fever Naples phlebovirus* species cluster, based on the L segment polymerase coding sequence. Gordil virus is used as the outgroup on the basis of the larger neighbour-joining tree (see Supplementary Figure 1). Comparable maximum parsimony trees for the M and L segments are given in Supplementary Figures 4 & 5.

### 3.2 Bayesian phylogeographic analysis of Toscana virus

TosV genotype C sequences from Croatia and Greece are only available for the L polymerase and S nucleoprotein coding sequences (compare Supplementary Tables 1 to 4). Therefore the question of the historical spread of the three clades focusses initially on the L polymerase and S nucleoprotein coding sequence datasets. Figure 2 shows the Bayesian phylogeographic tree for the L polymerase coding sequences, produced in BEAST. The location of the ancestor of the entire set of L sequences is undetermined, with a 0.56 probability of Tunisia versus Croatia or Greece. Genotype A spread initially within north Africa – Algerian and Tunisian sequences are the outgroups of genotype A – followed by a wider spread across the Mediterranean, to Turkey, Cyprus (possibly by way of Turkey, although with no statistical confidence), France and Italy. There may have been more than one introduction to France and Italy. Genotype B appears to have also spread within north Africa – into Morocco – and also across the Mediterranean to France and Spain. Genotype B may have been introduced to France and Spain on a single occasion. Genotype B subsequently spread to Turkey from north Africa.

**Figure 2.**
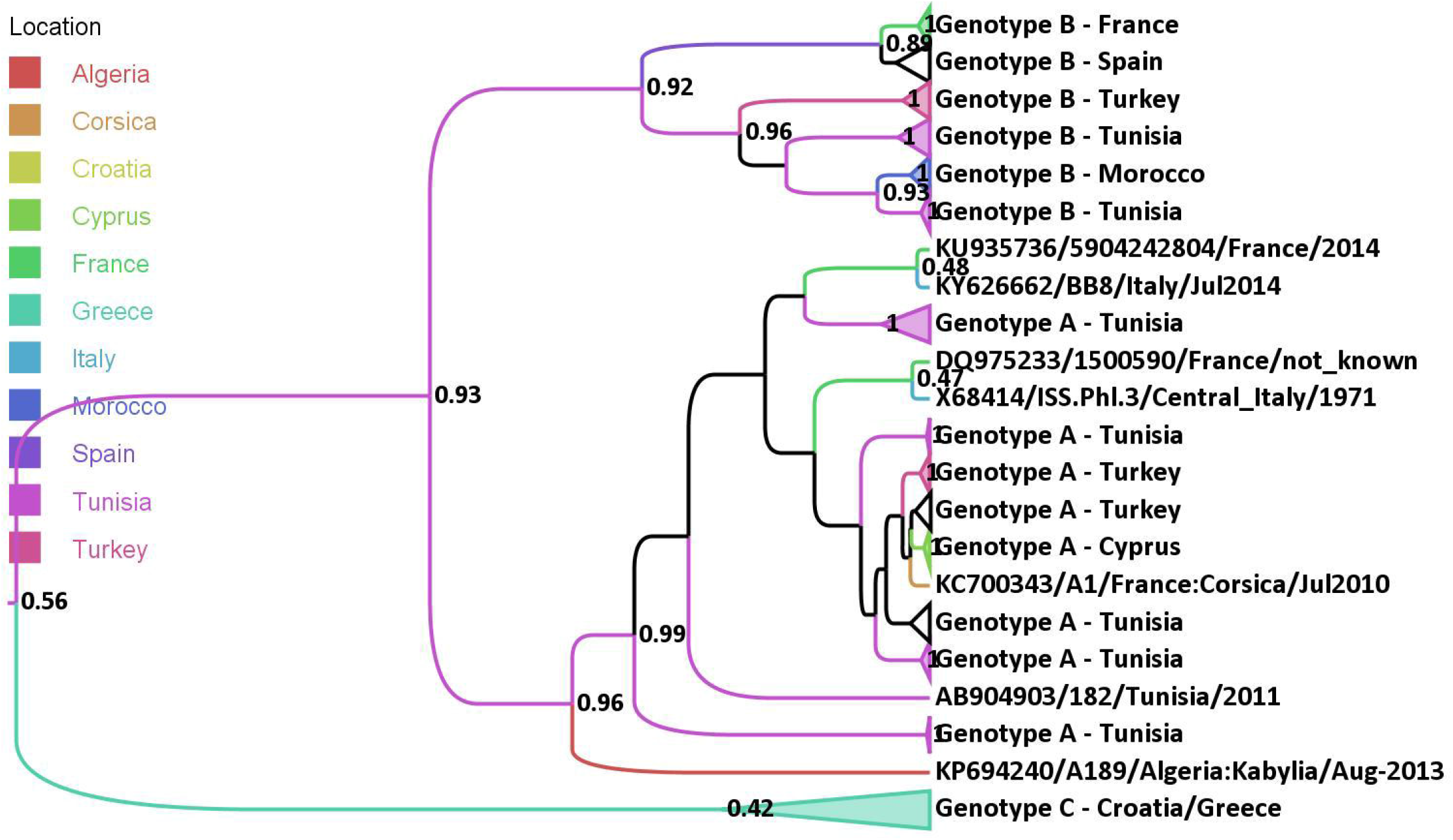
Bayesian phylogeographic tree for the L segment polymerase coding sequence, drawn in FigTree (Rambaut, 2014). No scale is given, as undated sequences are included and therefore no molecular clock calibration was performed. The node values are the location posterior probabilities. Branches are colour coded for most likely geographical location, according to the legend. Black branches are those for which posterior probability fell to a level where no phylogeographic reconstruction can be attempted. Where there are clades consisting entirely of one genotype and one geographical location, they are collapsed into triangles.

The phylogeographic analysis in Figure 2 is compromised by the short length of some of the L segment sequences used. Apart from the Italian, French and Spanish sequences, most are of a few hundred bases only. There are also comparatively few Italian L segment sequences in genotype A compared to the much larger diversity found in the M segment.

Figure 3 shows the equivalent phylogeographic tree for the nucleoprotein (N) coding sequence in the S segment. Most of these are full length sequences, by contrast with the predominantly fragmentary sequences for the L segment, although the N gene is the shortest coding sequence in TosV at 759 bases. The posterior probabilities for the geographic locations of the nodes, are therefore lower in Figure 3 than in Figure 2. As in the L segment, genotype C is the outgroup. Again, north African sequences occupy the basal positions within genotype A with subsequent spread into Europe and Turkey. A Portuguese sequence lies in the basal position for genotype B, but its corresponding L segment is not available, nor are any north African genotype B sequences.

**Figure 3.**
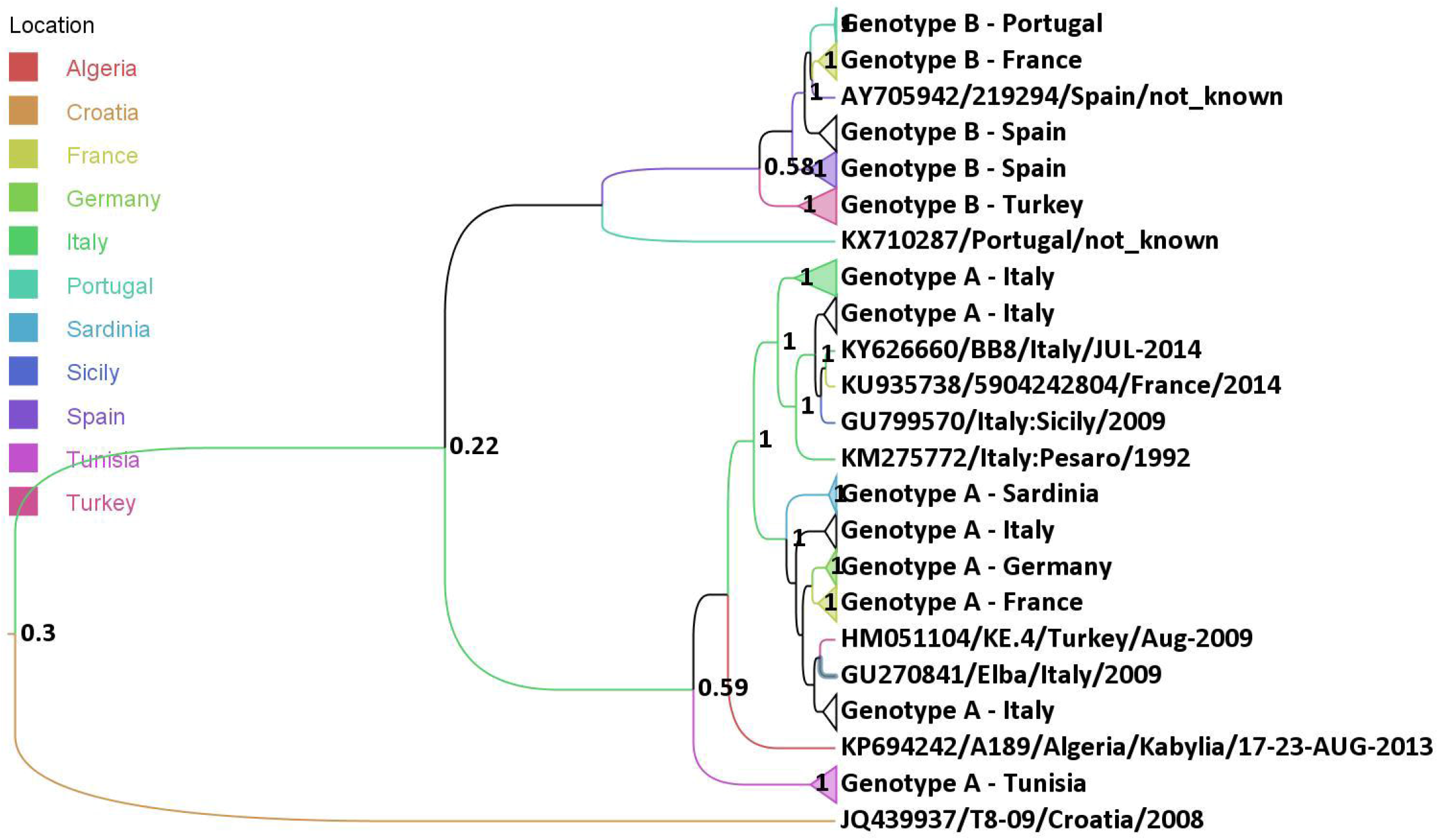
Bayesian phylogeographic tree for the S segment nucleoprotein coding sequence, drawn in FigTree (Rambaut, 2014). No scale is given, as undated sequences are included and therefore no molecular clock calibration was performed. The node values are the location posterior probabilities, colour coded according to the legend. Black branches are those for which posterior probability fell to a level where no phylogeographic reconstruction can be attempted. Where there are clades consisting entirely of one genotype and one geographical location, they are collapsed into triangles.

Supplementary Figures 6-7 show the phylogeographic trees for the M segment glycoprotein and the S segment NS coding sequences respectively. Although there are many more M segment sequences of genotype from Italy than there are L segment sequences, there are only a small number of M segment sequences from north Africa. M segment genotype B sequences are limited to France and Spain and there are no genotype C sequences. A similar situation prevails for S segment NS sequences.

### 3.3 Bayesian phylogenetic analysis of Toscana virus

A smaller set of sequences of greater length, and with sampling dates, was used to derive the timescale of TosV divergence. Table 1 shows the times of the most recent common ancestor (tMRCA) in years before the present (yBP) for each sequence set with the 95% HPDs in brackets.

**Table 1:**
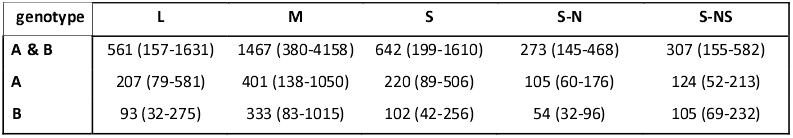
tMRCA values in years before present (yBP) with 95% highest posterior density (HPDs) in brackets for each segment. The S segment was analysed as a single alignment and then the component N and NS coding sequences were analysed in isolation. The “present” is the date of the most recently sampled sequence, 2015 in all cases except for M, which has one sequence from 2016. **A & B:** the tMRCA of TosV genotypes A and B. **A:** the tMRCA of TosV genotype A. **B:** the tMRCA of TosV genotype B. **L**: L segment genome. **M:** M segment genome. **S:** S segment genome. **S-N:** Nucleoprotein coding sequence. **S-NS:** Non-structural protein coding sequence.

Clock-like behaviour is not readily apparent in TosV genome evolution, and 95% HPDs are therefore wide, in some cases more than an order of magnitude. The common ancestor of the A and B genotypes, which is postulated based on Figure 2 to have existed in north Africa, is dated to several hundred years ago. The arrival of genotype A in Europe is dated slightly earlier than the arrival of genotype B, the former being in the order of hundreds of years ago and the latter in the order of decades.

For the M segment, Table 2 shows the tMRCAs of sub-genotype origins derived in the present study, compared with previous estimates (Zehender et al., 2009). Table 2 shows that the use of a restrictive prior on the substitution rate has not improved the precision of the 95% HPD ranges for the M segment as compared to a previous study (Zehender et al., 2009) where tip-date calibration only was used. The mean node tMRCA values are generally higher in the present study, although the mean tMRCAs are within 40 years of the previous estimates for 5 out of 7 of the nodes estimated. The exceptions are the MRCA of G2 and its parent node the MRCA of G1+2. One possible reason for this is that Zehender *et al* (Zehender et al., 2009) only analysed Italian sequences. However, our phylogeographic tree for the M segment (Supplementary Figure 6) shows that G1+2 does not constitute a monophyletic clade in Italy, but also includes Algerian and Tunisian strains. Because the relaxed exponential clock is the best model for TosV molecular evolution, and this model allows considerable rate variation between branches, the absence of the African branches may have distorted the mean substitution rate and hence the tMRCAs. We also note that Zehender *et al* (Zehender et al., 2009) used a relaxed lognormal clock rather than the relaxed exponential clock used here.

**Table 2:** tMRCA values for the M segment of TosV in years for before present (yBP) with 95% HPDs in brackets for each segment. The “present” is 2016. Sub-genotypes G1-4 are as previously defined by Zehender *et al* (Zehender et al., 2009). N.d.: not determined. **A & B:** the tMRCA of TosV genotypes A and B. **A:** the tMRCA of TosV genotype A. **B:** the tMRCA of TosV genotype B. **A-G1+2:** the tMRCA of TosV sub-genotypes G1 and G2. **A-G3+4:** the tMRCA of TosV sub-genotypes G3 and G4. **A – G1 to 4:** the tMRCA of TosV sub-genotypes G1 to G4 respectively.

| genotype | this study | Zehender et al.<br>(2009) |
| --- | --- | --- |
| **A & B** | 1467 (380-4158) | n.d. |
| **A** | 401 (138-1050) | 376 (75-856) |
| **B** | 333 (83-1015) | n.d. |
| **A - G1+2** | 289 (n.d) | 205 (40-485) |
| **A - G1** | 93 (54-194) | 55 (n.d.) |
| **A - G2** | 153 (58-396) | 51 (18-125) |
| **A - G3+4** | 142 (67-320) | 124 (35-252) |
| **A - G3** | 91 (46-201) | 69 (n.d.) |
| **A - G4** | 118 (60-256) | 82 (30-166) |

### 3.4 Natural selection on Toscana virus

Table 3 compares the substitution rate between the different segments of TosV and also across the three codon positions, as calculated by BEAST. The third codon position is less constrained by a factor of 2.2 to 2.7 relative to the overall substitution rate in all cases, and the ratio of non-synonymous to synonymous substitutions (dN/dS), calculated in Slr, ranges from 0.12 in the M segment coding sequence and the NS coding sequence of the S segment down to 0.02 in the N coding sequence of the S segment. Our value of 0.12 for the M segment is close to the 0.18 previously reported (Venturi et al., 2007). Such moderate to strong levels of overall selective constraint are suggestive of a virus that is well adapted to its hosts – both sandfly and human.

**Table 3:** The mean substitution rates in units of 10^−4^ substitutions per site per year for the segments of Toscana virus. The S segment overall rate is given as well as the rates for the two individual coding sequences. 95% HPD ranges are also given. **CP1-3:** the relative rates in the three coding positions compared to the overall rate. **Kappa:** the transition/transversion bias. **dN/dS:** omega (ratio of non-synonymous to synonymous substitution) N/A: not applicable. N/D: not determined.

| segment | mean | 95% HPD | CP1 | CP2 | CP3 | dN/dS | kappa |
| --- | --- | --- | --- | --- | --- | --- | --- |
| **L** | 2.38 | 0.85-3.99 | 0.25 | 0.11 | 2.64 | 0.04 | 4.39 |
| **M** | 1.08 | 0.28-1.92 | 0.5 | 0.27 | 2.23 | 0.12 | 6.16 |
| **S** | 2.23 | 0.97-3.49 | N/A | N/A | N/A | N/A | N/D |
| **S-N** | 2.94 | 1.69-4.26 | 0.08 | 0.14 | 2.78 | 0.02 | 6.88 |
| **S-NS** | 2.33 | 1.07-3.57 | 0.42 | 0.33 | 2.25 | 0.12 | 6.87 |

Even when the overall selective pressure is towards constraint, it is still possible for individual amino acid residues to be under positive selection. This was investigated using Slr (Massingham and Goldman, 2005). No residues in any of the segments were found to be positively selected at *p* < 0.05. Furthermore the N coding sequence of the S segment had no residues with sitewise dN/dS of >0.61. The residue with the highest sitewise dN/dS is residue 338 of the M segment coding sequence, as previously identified (Venturi et al., 2007). In the present study residue 338 has a sitewise dN/dS of 6.19, but the corrected significance value is *p* < 0.18 so it cannot be considered statistically significant. This residue presents a lysine/glutamate polymorphism with GAG, GAA and AAG codons all present. Substitution at this residue occurs sporadically throughout both A and B genotypes, suggesting that toggling of lysine with glutamate is occurring. Full Slr outputs are downloadable from http://dx.doi.org/10.17635/lancaster/researchdata/185.

### 3.5 Reassortment of genome segments in Toscana virus evolution

The simplest method of detection of genome reassortment is to look for incongruities in the positions of a given strain in phylogenetic trees from different segments. This can only be performed when more than one segment sequence per strain is available for a given strain. Supplementary Table 5 shows that 43 strains fall into this category, and that three Turkish strains (Thr2012, Ank2012 and pu3) are candidate reassortants with L segments of genotype A and S segments of genotype B. However, this must be treated with caution, since the Turkish sequences concerned were derived in two separate studies (Dincer et al., 2016; Ergunay et al., 2015) and although the authors noted the genotype in each case to be the same as that given in Supplementary Table 5, no mention was made of reassortment. It is not clear from the accompanying papers if the strain designation really indicates that each is derived from a single individual. Opportunities for reassortment can only occur where more than one genotype is circulating in the same area. Genotype C is so far confined to the Balkans, and all Italian and Spanish strains isolated so far are genotypes A and B respectively. The single strain isolated from Algeria is genotype A and the limited selection of Moroccan sequences are genotype B. Co-circulation of A and B occurs in France, Portugal, Tunisia and Turkey, so those countries would be the most likely locations for any reassortments to arise. This may constitute some circumstantial evidence for considering strains Thr2012, Ank2012 and pu3 as potential reassortants.

## 4. Discussion

### 4.1 The relationship of Toscana virus to other phleboviruses

The most parsimonious relationship between TosV and other members of the *Sandfly fever Naples phlebovirus* species group (SFNV) is shown in Figure 1 and Supplementary Figures 4 & 5. For the L segment (Figure 1), the Zerdali/Tehran virus pair are the most parsimonious sister clade of TosV with 70% bootstrap support. Moving outwards, the next sister clade at 71% bootstrap support is a quartet of Punique/Granada/Arrabida/Massilia viruses. The bootstrap confidence for that quartet clade is 98%. Figure 1 roots the tree on Gordil virus, following neighbour joining guide trees derived from a larger *Phlebovirus* set (Supplementary Figure 1). The reference SFNV L segment genome and that of two strains classed as SFNV-like constitute a clade with bootstrap confidences of 100%, and are found between the Gordil virus root and the in-group of 7 sequences. For the M segment, Sandfly fever Naples virus forms a clade with TosV and Tehran/Zerdali viruses (Supplementary Figure 4). Otherwise the phylogenetic relationships remain the same. By contrast for the S segment, the relative positions of the reference SFNV strains and the Zerdali/Tehran virus pair are reversed (Supplementary Figure 5). Despite the variation in bootstrap quality between the maximum parsimony trees for the three segments, their overall topological similarity suggests that TosV is part of a cluster of viruses that have evolved from an ancestral TosV/Tehran-Zerdali/Sandfly fever progenitor virus. The fact that topology is not exactly congruent between the three segments allows for the possibility that reassortment may have occurred between ancestral members of this species group.

Other phylogenetic analyses of the genus *Phlebovirus* appear in the literature. However, some of these do not use full length genome sequences (Ayhan et al., 2017; Punda-Polic et al., 2012), use a smaller set of SFNV species group viruses (Alkan et al., 2013; Collao et al., 2009; Depaquit et al., 2010; Punda-Polic et al., 2012), or use neighbour joining trees (Alkan et al., 2015; Ayhan et al., 2017), Bayesian trees (Punda-Polic et al., 2012) or UPGMA trees (Collao et al., 2009) rather than maximum parsimony. This paper is the first to present SFNV species group taxonomy using the full set of complete genome segments and using maximum parsimony tree building techniques.

### 4.2 Reassortment between Toscana virus genome segments

TosV is a segmented RNA virus, and both reassortment and recombination may theoretically occur in patients infected simultaneously with more than one strain. However, reassortment has only been demonstrated within the *Sandfly fever Naples virus phlebovirus* (SFNV) species cluster by Granada virus (Palacios et al., 2014) (see Figure 1 for phylogenetic relationships within SFNV) although laboratory experiments have shown the possibility of reassortment between TosV and Rift Valley fever virus (Accardi et al., 2001). Recombination has not been shown at all within SFNV but elsewhere in the genus *Phlebovirus* has been demonstrated in Severe Fever with Thrombocytopaenia Syndrome virus (SFTSV) (Shi et al., 2017). Given the previous lack of evidence for recombination in TosV, in this study we only search for signs of reassortment and leave recombination uninvestigated. The evidence for reassortment during TosV evolution is confined to a few Turkish strains (Supplementary Table 5). Although Turkey is one of the areas where genotypes A and B co-circulate, and thus a plausible location for the occurrence of TosV genotype co-infection with consequent reassortment, the evidence is not entirely conclusive based on the available sequences, their annotation and the lack of comment on the issue in the associated papers.

### 4.3 Route of spread of Toscana virus

The Bayesian phylogegraphic trees (Figures 2 and 3, Supplementary Figures 6 and 7) permit a reconstruction of the route of dispersal of TosV through the Mediterranean. Genotype C is the outgroup in the L segment and S-N trees. Figure 2 shows that the posterior probability for an origin of TosV in the Balkans versus north Africa is almost equally divided, with north Africa marginally favoured at 0.56. Therefore we postulate two alternative scenarios, both of which may be illustrated using Figure 4:

**Figure 4.**
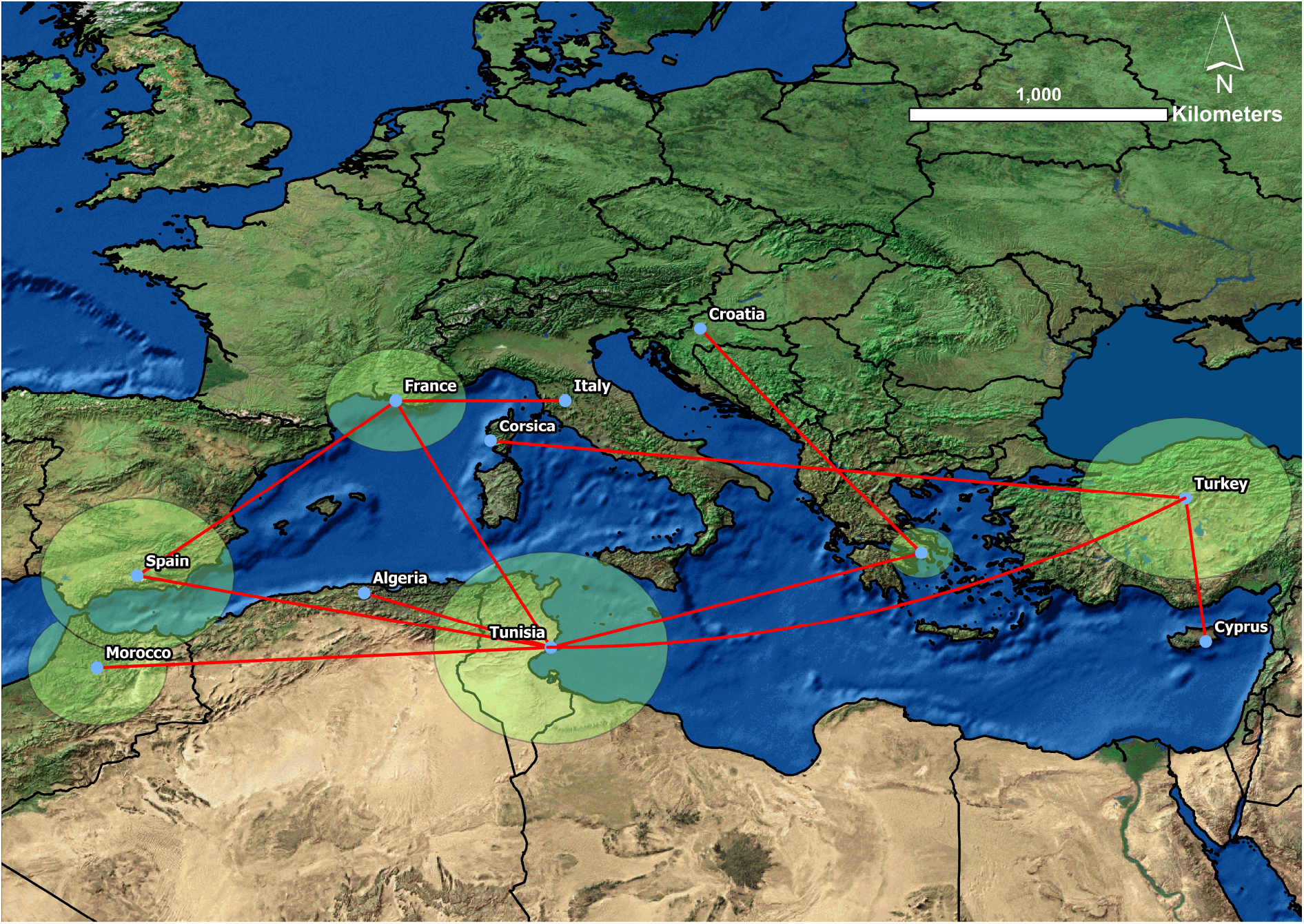
Bayesian phylogeographic tree for the L segment coding sequence (Figure 2) superposed on a map of the Mediterranean using SPREAD v.1.0.6. (Bielejec et al., 2011). The width of the circles indicate the number of sequences from that location. Since Figure 2 gives almost equal posterior probability to the origin of Toscana virus being located in the Balkans or in north Africa, this geographical network could be interpreted either with Tunisia or Greece as its origin.

1. TosV originated in north Africa and spread initially to the Balkans, giving rise to the ancestor of genotype C. Further diversification took place in Africa, possibly in Tunisia, producing ancestral forms of genotypes A and B. These spread in a complex manner into Europe and Turkey.
2. TosV originated in the Balkans, but then spread into north Africa where diversification into genotypes A and B occurred, prior to spreading into Europe and Turkey. The ancestral TosV subsequently evolved into modern genotype C in the Balkans.

Figure 4 may be interpreted in both ways, as the point of origin is not specified. Tunisia is either the point of origin of the common ancestor of all three TosV genotypes, or the first port of call after the point of origin in the Balkans. Although the scenarios differ as to the ultimate point of origin of TosV, both require north Africa to have been an important transmission hub into the western Mediterranean.

The L segment Bayesian phylogeographic tree (Figure 2) suggests two independent introductions to Europe of genotype A, each of which then diverged into Italy and France. Introduction 1 would have been the ancestor of genotype A strains BB8 (Italy) and 5904242804 (France), and introduction 2 the ancestor of strains ISS.Phl.3 (Italy) and 1500590 (France). The Corsican strain, A1, appears to be part of a Turkish/Cypriot cluster, so may represent a third introduction to France, but via the eastern Mediterranean rather than directly from north Africa (see also Figure 4). Genotype B, meanwhile, had a single introduction to Europe, which then diverged into French and Spanish varieties. Both genotypes A and B were introduced into Turkey from north Africa, after which genotype A was transmitted from Turkey to Cyprus and also to Corsica (Figures 1 and 4).

The S segment N coding sequence tree (Figure 3) supports this scenario while introducing some caveats. Once again, north African sequences occupy the basal position in genotype A. However, there are fewer genotype A Tunisian sequences for this segment, and therefore no means to corroborate the theory of more than one introduction of genotype A to Europe. By contrast, Italy is given as the most likely starting point of European genotype A diversification with posterior probability of 1. Once within Europe, genotype A spread from continental Italy to the islands of Sardinia and Elba, and also to France on two occasions. The first of these is associated with the strains BB8 (Italy) and 5904242804 (France), also paired in Figure 2, and the second of which involves strain 1500590 (France) with onward spread to Germany. The scenario for genotype B also becomes slightly more complicated once the information from Figure 3 is considered. A Portuguese strain is basal to genotype B suggesting, without good statistical confidence, that Portugal may have been the point of entry of genotype B to Europe, spreading from there into Spain and finally France. The presence of some Turkish genotype B sequences suggests a long-range spread eastwards. This is the second example (the first being the possible introduction of strain A1 from Turkey/Cyprus to Corsica, see Figures 1 and 4) of long-range spread superimposed on an underlying picture of shorter range geographical diffusion.

The M segment and S segment NS coding sequence trees (Supplementary Figures 6 and 7) are congruent with the scenario above. Those sequence sets contain less geographical diversity but nevertheless Supplementary Figure 7 supports the hypothesis, with 0.94 posterior probability, for the entry of genotype B to France from Spain. The Portuguese sequences are internal to the genotype B tree for segment S-NS but as in Figure 3, there is no statistical confidence for their placement.

### 4.4 Historical scenarios for the spread of Toscana virus

Although phleboviruses are found in both sandflies and the mammals upon which they feed, the long-range movement of insects is limited. The dispersal of Toscana virus is most probably a reflection of the spread of infected humans to new locations where ongoing transmission to local sandflies can occur, and new epidemic foci be established. The Mediterranean Sea has been a busy thoroughfare of trade, migration and warfare for thousands of years and there are many situations in which such dispersals could have occurred. The difficulty in dating the tMRCAs of TosV and its component genotypes (Table 1) means that it is largely a matter of speculation to associate TosV dispersal with specific historical events. The impossibility of deciding between the alternative Balkan and north African origins of TosV (Figures 1 and 4) introduces a further element of uncertainty.

Table 1 may be summarised approximately to show that the tMRCA of TosV existed around 380-468 yBP as this interval lies within the 95% HPDs of all segments. This is equivalent to a calendar date of 1547-1635 AD. This was a period during which the Turkish Ottoman Empire directly ruled the Balkans and controlled satellite states in north Africa, both of the candidate locations for the origin of TosV, providing numerous opportunities for the exchange of phleboviruses via administrative and commercial contacts. Similarly, a tMRCA range of 1839-1877 AD for genotype A and 1919-1932 AD for genotype B may be derived from the overlap in the 95% HPDs in Table 1. The French conquest of Algeria in 1830 and Tunisia in 1881, and the Italian conquest of Libya in 1911 may have increased opportunities for genotype A strains to enter Europe. For genotype B, the establishment of the Spanish protectorate in Morocco in 1912 may have performed the same function. Alternative scenarios may be derived.

Figure 1 suggests that TosV may have Tehran (Iran) and Zerdali (Turkey) viruses as its nearest relatives. This also provides some circumstantial evidence that the deeper history of TosV, prior to the most recent common ancestor of the extant genotypes, may have been within the boundaries of the Ottoman and Persian Empires in more ancient times.

### 4.5 Selection and adaptation in Toscana virus

Slr and BEAST analysis (Table 3) support the previous calculation of moderate constraining selective pressure on all segments of TosV. Additionally, the candidate positively selected site in the glycoprotein at residue 338 (Venturi et al., 2007) has a less statistically significant signal in the current study than previously. The overall picture is of a virus that is well adapted to its host. The absence of the typical immune evasion strategy pattern of strong positive selection on a small number of epitope residues, suggests that this scenario does not apply to TosV. The several centuries of TosV presence in humans in Europe, Asia and Africa may have provided opportunity for functional adaptation. Since TosV’s closest relatives are also viruses of humans (Figure 1), this adaptational process may be part of a much longer evolutionary adaptation to human hosts of phleboviruses in general.

### 4.6 Improving the resolution of the historical scenario of Toscana virus spread

Calibration of molecular clocks for TosV has remained a challenging problem, as in previous studies. Geographical information is patchy, with some countries (e.g. Tunisia) delivering a wealth of short sequences but no full length genomes, and others (e.g. Italy) with many more sequences of one segment over the others. Many other probable areas for TosV circulation in the Mediterranean basin have no sequence coverage at all. Future work requires the generation of full genome sequences for those north African strains for which only partial sequences are currently available. More L segment sequences from Italy are needed, and more sequences of all kinds are required from genotype C in the Balkans.

## Author statements

### Funding Information

None.

## Acknowledgments

We thank Dr Tim Massingham and Dr Nick Goldman (European Bioinformatics Institute, Cambridge, UK) for providing access to Slr 1.5.0 prior to release, and Dr Luigi Sedda (Lancaster University) for assistance with production of Figure 4. DG thanks Dr Rod Dillon and Prof Paul Bates (Lancaster University) for introducing him to the world of sandfly-related diseases.

## Conflicts of interest

None.

## Ethical statement

Not applicable.

## Data availability statement

Raw data and intermediate analysis files are freely available at: http://dx.doi.org/10.17635/lancaster/researchdata/185.

